# Upregulation of the AMPK-FOXO1-PDK4 pathway is a primary mechanism of pyruvate dehydrogenase activity reduction and leads to increased glucose uptake in tafazzin-deficient cells

**DOI:** 10.1101/2024.02.03.578755

**Authors:** Zhuqing Liang, Tyler Ralph-Epps, Michael W. Schmidtke, Pablo Lazcano, Simone W. Denis, Mária Balážová, Mohamed Chakkour, Sanaa Hazime, Mindong Ren, Michael Schlame, Riekelt H. Houtkooper, Miriam L. Greenberg

**Affiliations:** Department of Biological Sciences, Wayne State University, Detroit, Michigan, United States; Laboratory Genetic Metabolic Diseases, Amsterdam UMC, University of Amsterdam, Amsterdam, The Netherlands; Amsterdam Gastroenterology Endocrinology and Metabolism institute, Amsterdam, The Netherlands; Amsterdam Cardiovascular Sciences institute, Amsterdam, The Netherlands; Emma Center for Personalized Medicine, Amsterdam UMC, Amsterdam, The Netherlands; Department of Membrane Biochemistry, Institute of Animal Biochemistry and Genetics, Centre of Biosciences, Slovak Academy of Sciences, 84005 Bratislava, Slovakia; Department of Anesthesiology, Perioperative Care, and Pain Medicine, Grossman School of Medicine, New York University, New York, New York, United States

**Author notes:** For correspondence: Miriam L. Greenberg.

**Keywords:** cardiolipin, pyruvate dehydrogenase complex, pyruvate dehydrogenase kinase, FOXO, AMP-activated kinase (AMPK), glucose uptake, GLUT4

## Abstract

Barth syndrome (BTHS) is a rare disorder caused by mutations in the *TAFAZZIN* gene. Previous studies from both patients and model systems have established metabolic dysregulation as a core component of BTHS pathology. In particular, features such as lactic acidosis, pyruvate dehydrogenase (PDH) deficiency, and aberrant fatty acid and glucose oxidation have been identified. However, the lack of a mechanistic understanding of what causes these conditions in the context of BTHS remains a significant knowledge gap, and this has hindered the development of effective therapeutic strategies for treating the associated metabolic problems.

In the current study, we utilized tafazzin-knockout C2C12 mouse myoblasts (TAZ-KO) and cardiac and skeletal muscle tissue from tafazzin-knockout mice to identify an upstream mechanism underlying impaired PDH activity in BTHS. This mechanism centers around robust upregulation of pyruvate dehydrogenase kinase 4 (PDK4), resulting from hyperactivation of AMP-activated protein kinase (AMPK) and subsequent transcriptional upregulation by forkhead box protein O1 (FOXO1). Upregulation of PDK4 in tafazzin-deficient cells causes direct phospho-inhibition of PDH activity accompanied by increased glucose uptake and elevated intracellular glucose concentration. Collectively, our findings provide a novel mechanistic framework whereby impaired tafazzin function ultimately results in robust PDK4 upregulation, leading to impaired PDH activity and likely linked to dysregulated metabolic substrate utilization. This mechanism may underlie previously reported findings of BTHS-associated metabolic dysregulation.

## Introduction

Barth syndrome (BTHS) is a rare genetic disorder caused by mutations in the *TAFAZZIN* gene, which encodes a transacylase responsible for remodeling the vital mitochondrial lipid cardiolipin. BTHS is life-threatening, as patients often suffer from a host of associated pathologies, including cardiomyopathy, skeletal myopathy, neutropenia, and various metabolic irregularities. Currently, there is a lack of effective therapies available for this disease, highlighting the need for a better understanding of the pathophysiology. Recent clinical studies have revealed a correlation between impaired cardiac bioenergetics, reduced skeletal muscle mass, and defective substrate metabolism in BTHS patients (Cade et al., 2019). In particular, during submaximal exercise, patients exhibit blunted glucose and fatty acid oxidation (FAO), increased glucose turnover rate, and higher concentrations of plasma lactate, whereas at rest, plasma fatty acid content is elevated while glucose concentrations are similar to healthy controls (Cade et al., 2013; Cade et al., 2019).

Although glucose oxidation and FAO represent alternative bioenergetic strategies, both pathways converge on acetyl-CoA production. Coordinate regulation of glucose oxidation and FAO is generally attributed to the Randle cycle. However, activation of AMP-activated protein kinase (AMPK) in response to metabolic stressors, such as reduced substrate availability, increased energy demand, or hypoxia, can also increase FAO and glucose oxidation independent of regulation by the Randle cycle (Carling et al., 2003). Acetyl-CoA levels are substantially reduced in CL-deficient yeast, and recent work has shown that tafazzin-knockout C2C12 mouse myoblasts (TAZ-KO) exhibit a robust reduction in acetyl-CoA resulting from reduced pyruvate dehydrogenase (PDH) activity (Raja et al., 2017; Li et al., 2019). PDH functions as a regulatory hub by governing the conversion of glycolysis-derived pyruvate into acetyl-CoA (Randle et al., 1963; Hue and Taegtmeyer, 2009). Inhibition of PDH leads to the preservation of pyruvate and lactate, whereas PDH activation facilitates glucose oxidation by producing glycolysis-derived acetyl-CoA to be utilized by the tricarboxylic acid (TCA) cycle and electron transport chain (ETC).

PDH activity is tightly regulated through reversible phosphorylation on the E1 subunit, which is modulated by the relative activities of pyruvate dehydrogenase kinases (PDK) and pyruvate dehydrogenase phosphatase. This intricate regulatory mechanism enables PDH to coordinate glucose oxidation, thereby maintaining metabolic flexibility in substrate selection. Disruption of this regulatory process has been implicated in various metabolic disorders, including diabetes, cancers, and heart failure (Sheeran et al., 2019), and this has prompted investigations into therapeutic interventions targeting PDH in conditions such as nonalcoholic fatty liver disease (Saed et al., 2021) and obesity (Al Batran et al., 2020).

Impaired PDH activity has been observed in TAZ-KO C2C12 myoblasts as well as in heart tissue from TAZ-KO mice and myocardial extracts from tafazzin-kockdown mice (Cade et al., 2019)(Li et al., 2019; Greenwell et al., 2021). Therefore, we hypothesize that dysregulation of PDH may contribute to the metabolic aberrancies in FAO and glucose utilization observed in BTHS patients. Despite these observations, the molecular mechanism underlying impaired PDH activity in BTHS remains largely unexplored, and this knowledge gap has precluded the development of effective therapeutic interventions for treating the metabolic component of BTHS. The current study utilized TAZ-KO C2C12 myoblasts and TAZ-KO mouse tissue to elucidate a key mechanism underlying PDH inhibition in the context of tafazzin deficiency and identify the associated alterations in glucose utilization resulting from dysregulation of PDH. Our data indicate that the absence of tafazzin leads to hyperactivation of the metabolic stress sensor AMPK, which subsequently acts through a downstream signaling cascade to trigger robust upregulation of PDK4, inhibition of PDH activity, increased glucose uptake, and intracellular glucose accumulation.

## Results

### PDK4 is upregulated in tafazzin-deficient cells

The PDH complex, which converts pyruvate to acetyl-CoA, plays a central role in coordinating metabolic substrate utilization and maintaining substrate flexibility (for review, see (Park et al., 2018)). Given its importance, aberrant suppression of PDH (i.e., PDH deficiency) is associated with numerous metabolic diseases, including BTHS (Li et al., 2019; Greenwell et al., 2021; Ghosh et al., 2022). However, the underlying cause of PDH deficiency in BTHS has remained unknown.

PDH is reversibly regulated by phosphorylation, with upregulation of pyruvate dehydrogenase kinase (PDK) and/or downregulation of pyruvate dehydrogenase phosphatase (PDP) acting as direct negative regulatory mechanisms for PDH activity. Therefore, to test whether inhibition of PDH in TAZ-KO cells results from altered expression of these phospho-regulatory enzymes, we assayed mRNA and protein expression of all PDK and PDP isoforms. Strikingly, this analysis revealed that mRNA levels of PDK4, the predominant isoform expressed in C2C12 cells (Fig. 1A), are increased substantially (2- to 6-fold) in both pre- and post-differentiation-induced TAZ-KO myoblasts as well as in cardiac and skeletal muscle from TAZ-KO mice (Fig. 1A, B). Western blot analysis showed that PDK4 protein is also increased in TAZ-KO myoblasts and cardiac muscle (Fig. 1C), suggesting that upregulation of PDK4 leads to decreased PDH activity in TAZ-KO cells.

**Figure 1.**
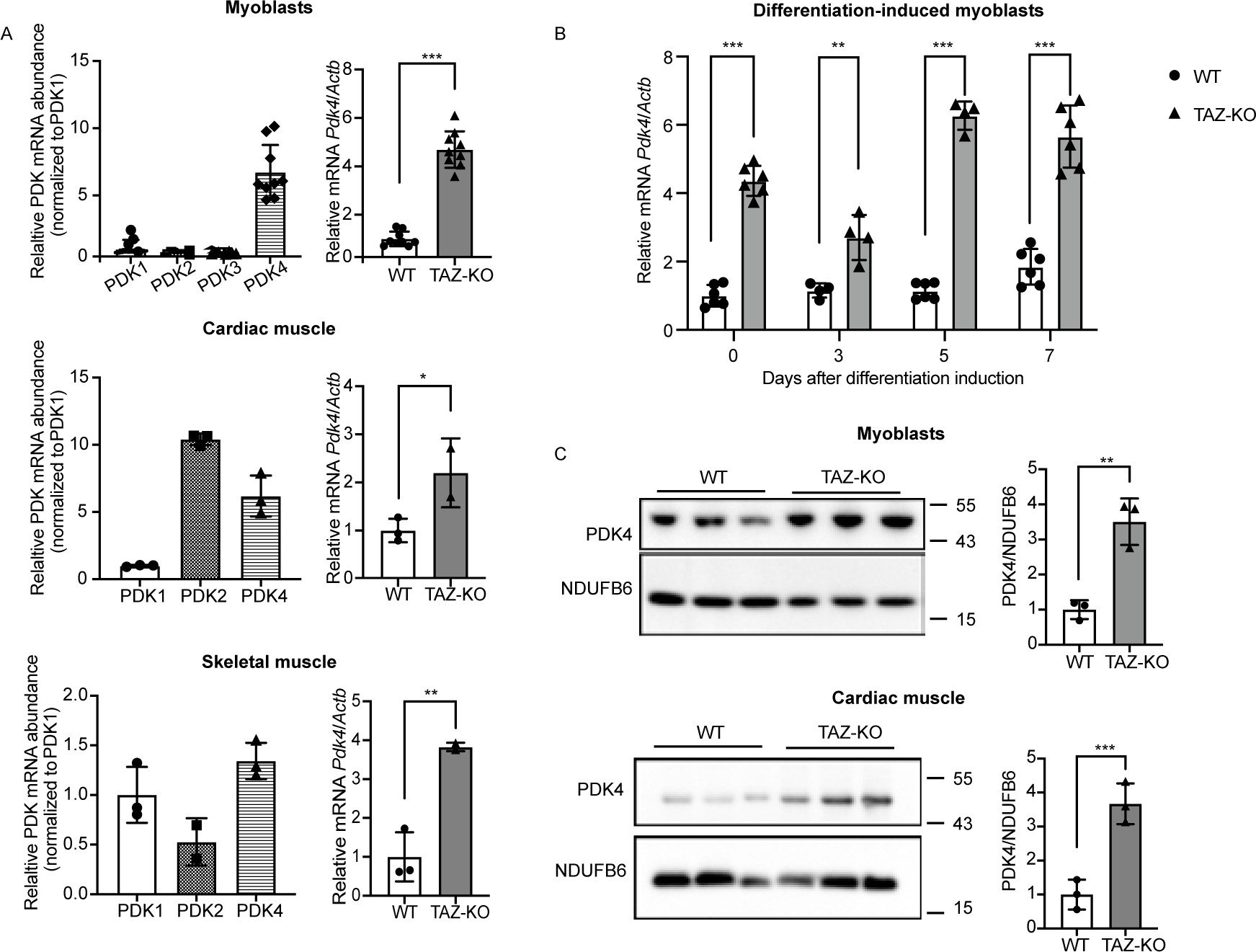
PDK4 is upregulated in tafazzin-deficient cells. PDK4 and other isoforms levels were evaluated using qPCR (A) (B) or Western blot (WB) analysis (C) of whole-cell lysates extracted from WT and TAZ-KO myoblasts, differentiation-induced myotubes, and mouse muscle tissue. The figures in panel (C) are representative images with each lane representing one biological replicate. The mRNA expression levels were normalized to the internal control *Actb,* and protein levels were normalized to NDUFB6. Densitometry was quantified using Image J and statistical significance was determined with an unpaired, two-tailed Student’s *t*-test. The data points represent mean ±S.D. (*error bars*) for each individual biological replicate of each group. * 0.05<*p*<0.1, ** 0.01<*p*<0.05, *** 0.001<*p*<0.01.

### PDK4 upregulation leads to PDH deficiency in TAZ-KO cells

To test whether the elevated PDK4 levels observed in TAZ-KO cells result in the previously reported phospho-inhibition of PDH (Li et al., 2019), we treated cells with the PDK inhibitor dichloroacetate (DCA). DCA treatment resulted in a reduction in PDK4 protein levels, as well as a concomitant decrease in PDH phosphorylation (Fig. 2A). Because inhibition of PDK by DCA is not specific to the PDK4 isoform, we also utilized the complementary approach of PDK4-targeted siRNA knockdown to test whether inhibition of PDK4 alone is sufficient to rescue the phospho-PDH phenotype. Our results consistently showed that PDH phosphorylation is reduced to WT levels in TAZ-KO cells treated with PDK4-targeted siRNA (Fig. 2B). As a metabolic hub, PDH activity can indirectly influence mitochondrial oxidative phosphorylation, and TAZ-KO myoblasts have been previously shown to exhibit defects in mitochondrial oxygen consumption rate (OCR) (Lou et al., 2018; Ji et al., 2022). Therefore, to determine whether inhibiting PDK4 improves mitochondrial function, we tested the ability of DCA treatment to restore OCR in TAZ-KO cells. Treating TAZ-KO cells with 5 mM DCA resulted in a slight increase in basal respiration and a more pronounced increase in maximal respiration (Fig. 2C). These findings strongly suggest that PDK4 upregulation is directly responsible for phospho-inhibition of the metabolic regulator PDH and that restoring PDH activity improves mitochondrial function in TAZ-KO cells.

**Figure 2.**
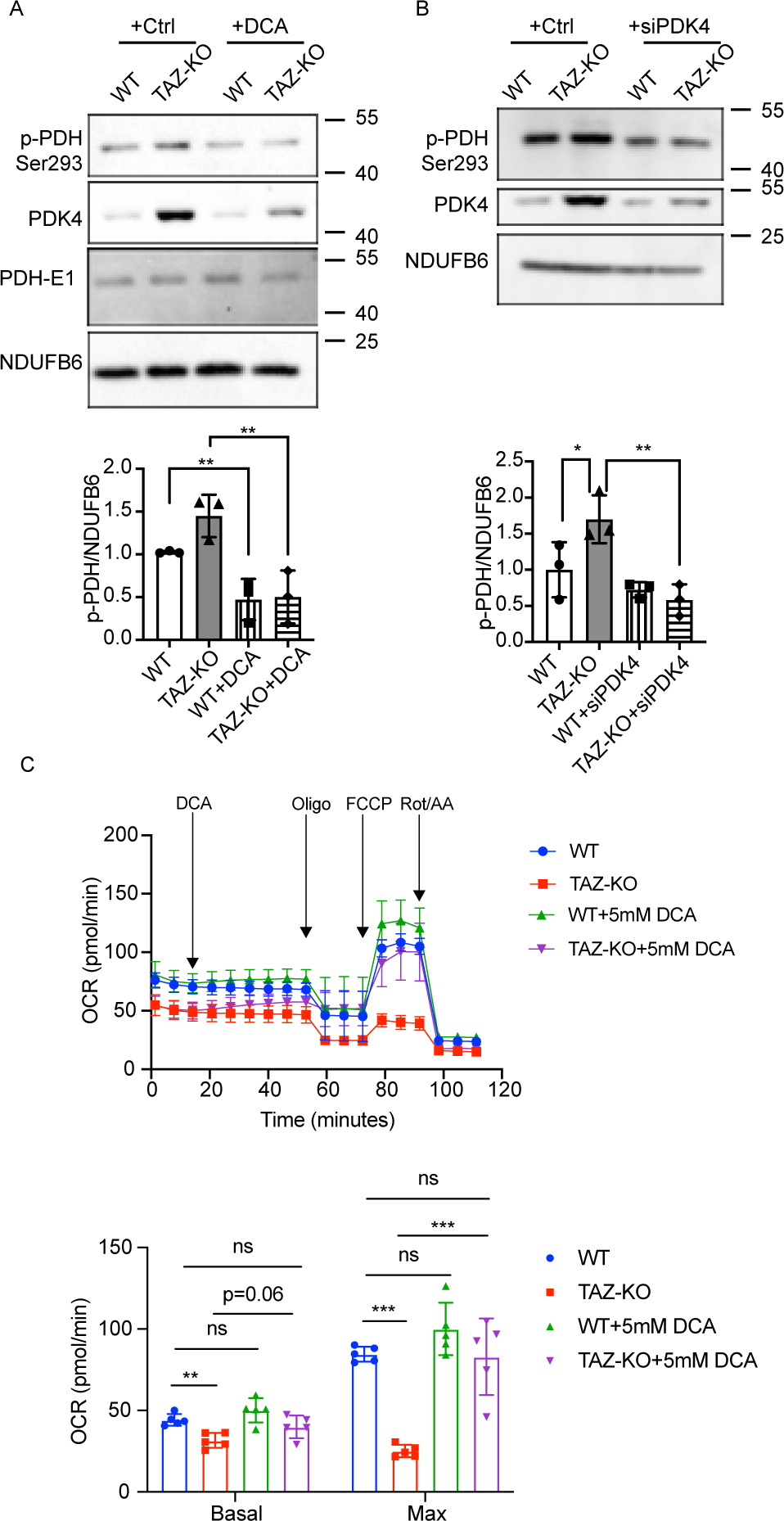
Inhibiting PDK4 decreases PDH phosphorylation in TAZ-KO cells. Myoblasts were treated with either 5 mM DCA for 24 h (A) or PDK4-targeted siRNA (B). Phosphorylation of PDH on Ser^293^ (p-PDH) was assessed via WB. Image J software was used for quantitative analysis of p-PDH (bottom panels). (C) Oxygen consumption rates (OCR) of myoblasts subjected to 5 mM DCA treatments. n=5. The data points represent mean ±S.D. (*error bars*) for each individual biological replicate of each group. * 0.05<*p*<0.1, ** 0.01<*p*<0.05, *** 0.001<*p*<0.01.

### Upregulation of FOXO1 activates PDK4 transcription in TAZ-KO cells

In light of our findings that collectively implicate PDK4 upregulation as a primary mechanism underlying PDH activity reduction in TAZ-KO cells, we next aimed to investigate the mechanism of PDK4 upregulation, with the goal of highlighting potential therapeutic targets in the underlying pathway. In skeletal muscle, PDK4 expression is enhanced by several transcription factors, including forkhead box protein O1 (FOXO1) (Furuyama et al., 2003). Dysregulation of FOXO1 has been previously associated with metabolic disease and muscle atrophy (Wu et al., 2008; Puthanveetil et al., 2013; Oyabu et al., 2022).

To determine whether upregulation of PDK4 in TAZ-KO cells can be attributed to increased FOXO1-mediated transcription, we used complementary approaches to probe FOXO1 activity. This analysis revealed an increase in both FOXO1 mRNA and protein levels in TAZ-KO cells, indicating increased *de novo* synthesis of FOXO1 (Fig. 3A), as well as enrichment of FOXO1 in the nuclei of TAZ-KO cells, suggesting enhanced transcriptional regulatory activity (Fig. 3B, C) (Wu et al., 2014). We also confirmed direct regulation of PDK4 by FOXO1 using chromatin immunoprecipitation (Fig. 3D) and noted an increase in the mRNA levels of other previously reported FOXO1 substrates, including *Cdkn1a*, *Gagarapl1*, *Gadd45*γ, and *Plk2* (Fig. 3E), suggesting an overall increase in FOXO1-mediated transcription in TAZ-KO cells. Consistent with the observations in TAZ-KO myoblasts, tissue from TAZ-KO mouse hearts also showed increased FOXO1 mRNA (Fig. 4A), a nearly significant increase in FOXO1 protein (Fig. 4B), and an increase in the mRNA of FOXO1-regulated transcriptional targets (Fig. 4C). Collectively, these findings suggest that upregulation and nuclear enrichment of FOXO1, observed in both the *in vitro* TAZ-KO myoblast model and *in vivo* TAZ-KO mouse cardiac tissue, directly increase transcription of *Pdk*4.

**Figure 3.**
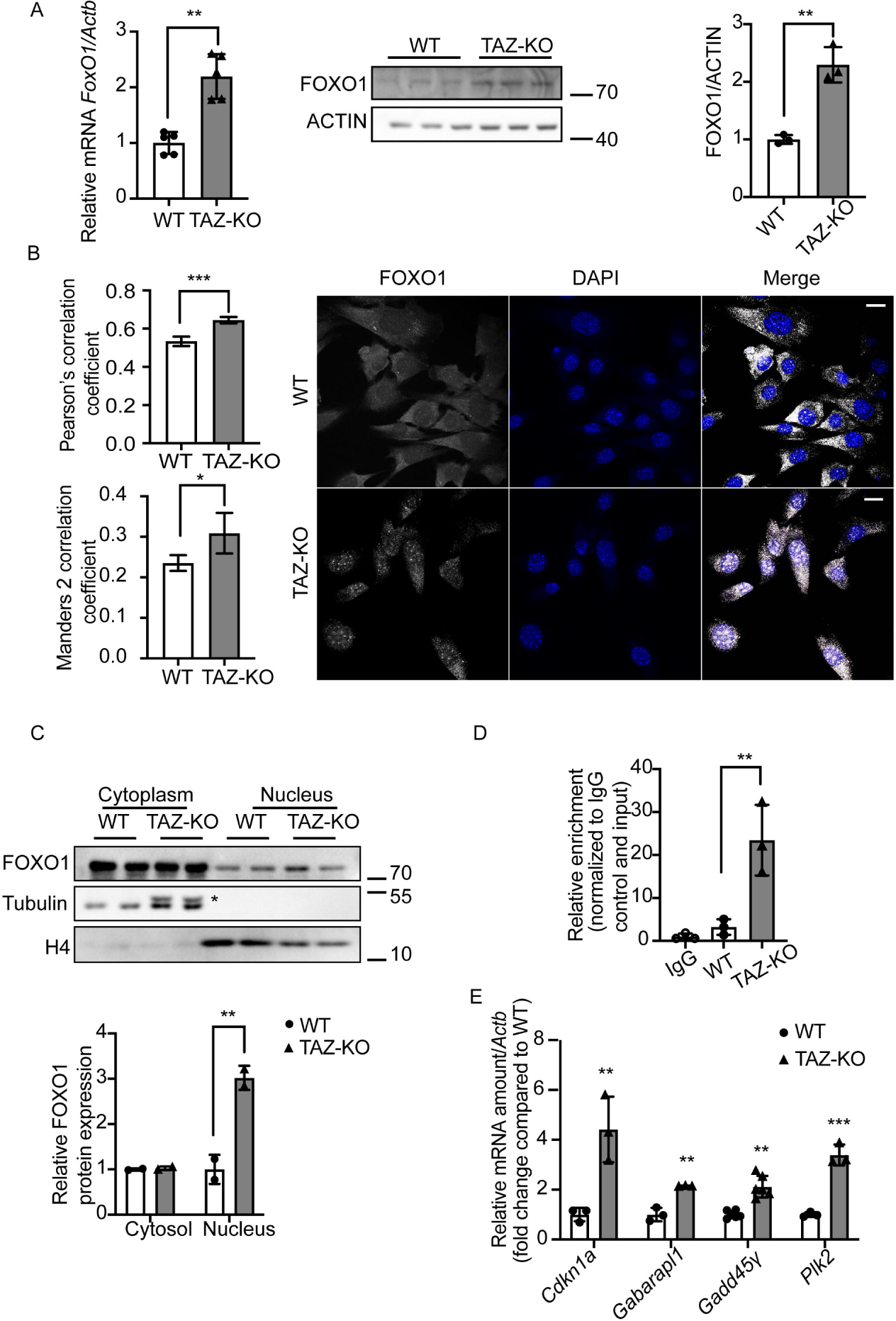
Upregulation of FOXO1 activates PDK4 transcription in TAZ-KO cells. (A) Total mRNA and whole-cell lysate were extracted from both WT and TAZ-KO myoblasts, and the mRNA and protein levels of FOXO1 were measured using real-time qPCR and WB analysis, respectively. Data are presented as fold-change relative to WT, and each lane represents an individual biological replicate. (B) *Right:* The subcellular localization of FOXO1 was detected through immunofluorescence (IF) confocal microscopy, with representative figures showing the relative distribution of FOXO1 in WT and TAZ-KO myoblasts. *Left:* Quantification of colocalization between FOXO1 and DAPI based on Pearson’s and Mander’s 2 colocalization analysis (Manders et al., 1992; Manders et al., 1993). The scale bar equals 20 µm. (C) Nuclear protein fractionation was performed, followed by WB analysis using tubulin and histone 4 (H4) as internal controls for cytoplasmic and nuclear fractions, respectively. * indicates PDK4 band residue after stripping the membrane. (D) Chromatin immunoprecipitation/qPCR was performed in myoblasts with either an IgG control or anti-FOXO1 antibody. The *Pdk4* promoter was detected using the specific primers listed in “Experimental procedures”. (E) Real-time qPCR was used to measure mRNA of the FOXO1 targets *Cdkn11* (Cyclin Dependent Kinase Inhibitor 1A), *Gabarapl1* (Gamma-aminobutyric acid receptor-associated protein-like 1), *Gadd45γ* (Growth Arrest and DNA Damage Inducible γ), and *Plk2* (Polo Like Kinase 2), using *Actb* as the internal control. Data points represent mean ±S.D. (*error bars*) for each individual biological replicate of each group. * 0.05<*p*<0.1, ** 0.01<*p*<0.05, *** 0.001<*p*<0.01.

**Figure 4.**
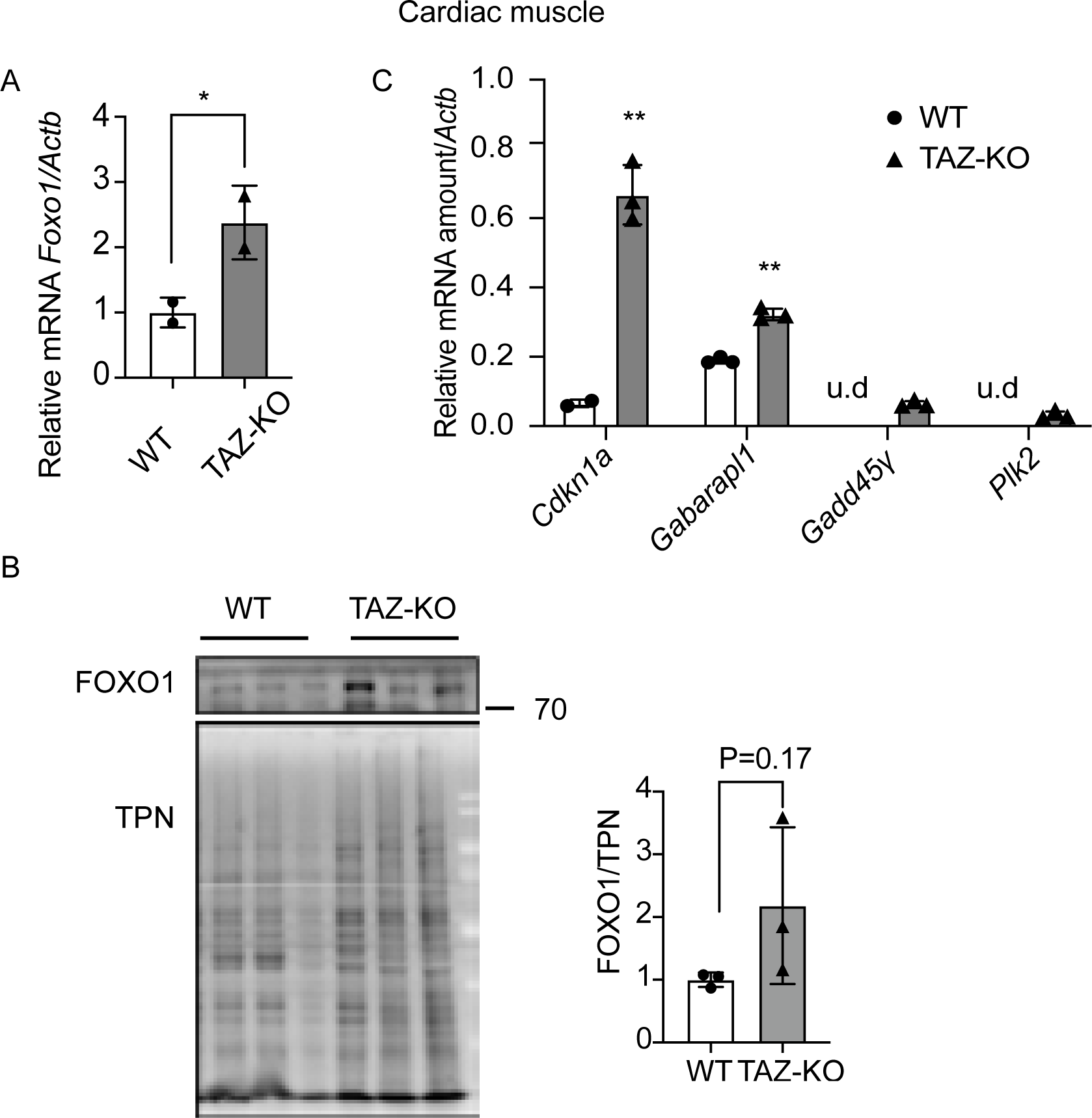
TAZ-KO cardiac muscle shows enrichment of nuclear FOXO1. Total mRNA and whole-cell lysate were extracted from both WT and TAZ-KO mouse cardiac tissues, and FOXO1 mRNA (A) and protein (B) were measured using real-time qPCR and WB analysis, respectively. A total protein stain was used for WB analysis normalization (TPN). Data are presented as fold-change relative to WT, with each lane representing an individual biological replicate. (C) Real-time qPCR was used to measure the mRNAs of four FOXO1 substrates (*Cdkn11, Gabarapl1, Gadd45γ, and Plk2*) using *Actb* as the internal control. Data points represent mean ±S.D. (*error bars*) for each individual biological replicate of each group. * 0.05<*p*<0.1, ** 0.01<*p*<0.05, *** 0.001<*p*<0.01.

### AMPK mediates the enrichment of nuclear FOXO1 and PDK4 upregulation in TAZ-KO cells

FOXO1 expression and nuclear translocation have been shown to be regulated by multiple mechanisms, including phosphorylation by the highly conserved cellular energy sensor AMPK (Yun et al., 2014). Activation of AMPK functions as a ‘metabolic switch’, shifting cellular metabolism away from ATP-consuming anabolic reactions towards ATP-producing catabolic processes, including FAO and glucose oxidation (Winder and Hardie, 1999; Samovski et al., 2015; Herzig and Shaw, 2018). The Randle cycle is generally attributed with balancing utilization of fatty acids versus glucose as metabolic substrates. However, in the face of metabolic stress, AMPK activation has been shown to increase both FAO and glucose oxidation independent of Randle cycle-mediated regulation (Carling et al., 2003).

Intriguingly, increased AMPK activity has been observed in BTHS patient lymphoblasts (Mejia et al., 2017) and tafazzin-knockdown neonatal ventricular myocytes (He, 2010), suggesting that hyperactivation of AMPK may play a causative role in the context of BTHS-associated metabolic dysregulation. Therefore, we hypothesized that the enrichment of nuclear FOXO1 in TAZ-KO cells is caused by activation of AMPK. Because AMPK is activated by phosphorylation on residue Thr^172^, we measured the levels of Thr^172^-phosphorylated AMPK (pAMPK) as an indicator of AMPK activity (Stein et al., 2000; Suter et al., 2006; Sanders et al., 2007). As predicted, levels of pAMPK were robustly increased in TAZ-KO cells (Fig. 5A), suggesting an increase in downstream AMPK activity.

**Figure 5.**
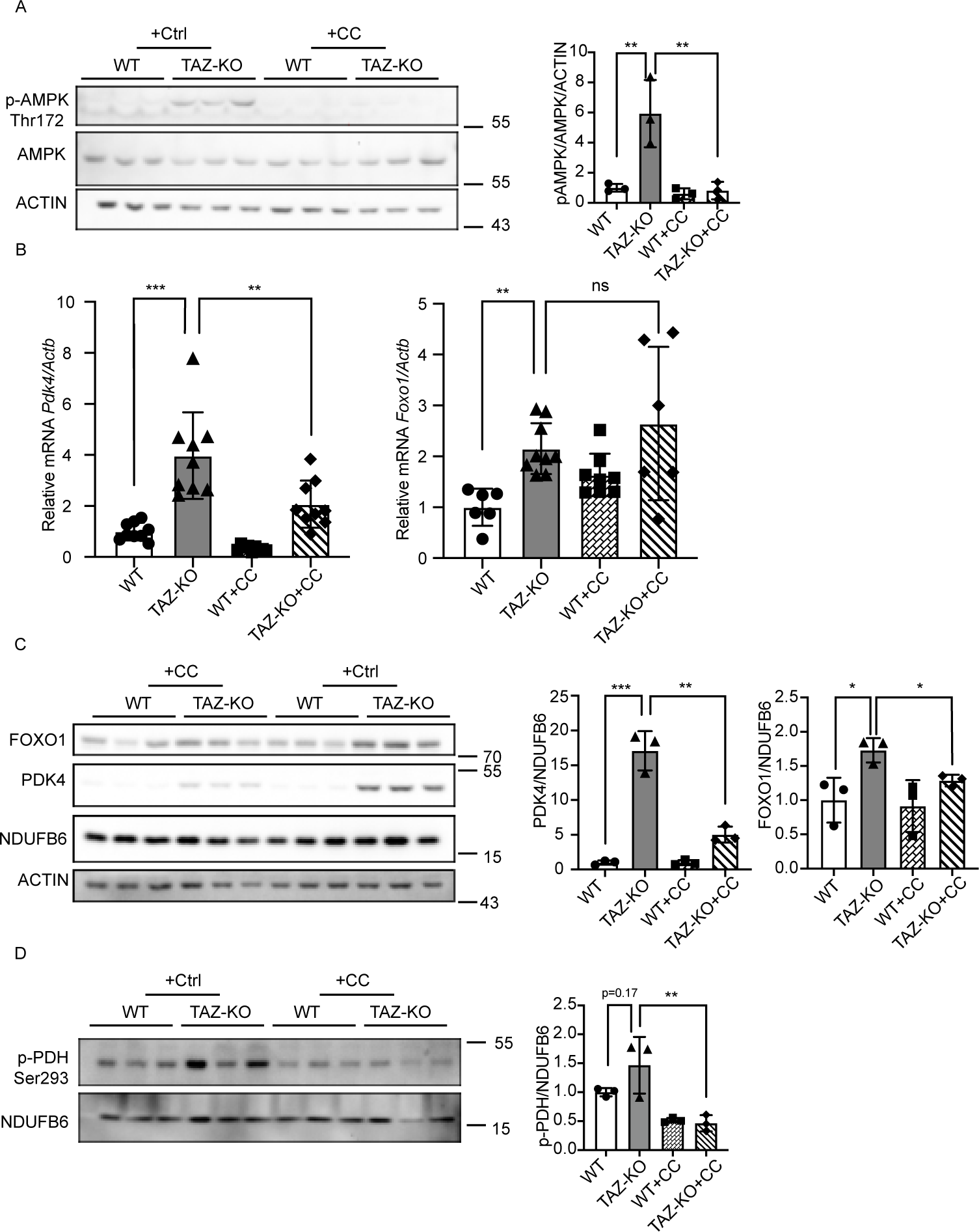
AMPK-mediated upregulation of FOXO1 increases PDK4 in TAZ-KO cells. (A) Phosphorylation of AMPK residue Thr^172^ (p-AMPK) was assayed in WT and TAZ-KO myoblasts treated with the AMPK inhibitor compound C (CC) or vehicle. 10 μg of total protein was loaded for each sample, and ACTIN was used as an internal control. Representative images and quantitative analysis are shown. (B) Total mRNA was extracted from myoblasts, and *Pdk4* and *Foxo1* mRNA levels were measured via real-time qPCR following treatment with CC (10 μM, 16 h) or vehicle. (C) Protein expression of FOXO1 and PDK4 in myoblasts was measured via WB analysis following treatment with CC (10 μM, 16 h) or vehicle. Representative images and corresponding quantitative analyses are shown. (D) In myoblasts, p-PDH was measured by WB analysis following treatment with CC (10 μM, 16 h) or vehicle. Representative images and corresponding quantitative analyses are shown. Data points represent mean ±S.D. (*error bars*) for each individual biological replicate of each group. * 0.05<*p*<0.1, ** 0.01<*p*<0.05, *** 0.001<*p*<0.01.

As a transcription factor, nuclear translocation of FOXO1 is an essential prerequisite for its role in mediating transcription, and previous work has demonstrated that phosphorylation of FOXO1 by pAMPK is sufficient to increase its nuclear enrichment and transcriptional activation of target genes (Yun et al., 2014). Therefore, we sought to assay the expression patterns of FOXO1 and PDK4 following treatment with the widely used AMPK inhibitor, compound C (CC) (Zhou et al., 2001). We found that treating TAZ-KO cells with CC (10 μM, 16 h) resulted in a significant decrease in AMPK activity (Fig. 5A), as well as a decrease in mRNA levels of PDK4, but not FOXO1 (Fig. 5B). However, at the protein level, AMPK inhibition led to a reduction in both PDK4 and FOXO1 (Fig. 5C), and a decrease in PDH phosphorylation relative to controls (Fig. 5D).

Given the modest effect of AMPK inhibition on FOXO1 mRNA and protein relative to PDK4, we reasoned that AMPK likely regulates FOXO1 by a predominantly post-translational mechanism to influence its subcellular localization. To test this prediction, we conducted immunocytochemistry of FOXO1 in WT and TAZ-KO cells with and without CC treatment and examined its colocalization with the nuclear marker DAPI. Our results showed that inhibition of AMPK in TAZ-KO cells reduced the nuclear localization of FOXO1 to WT levels (Fig. 6A). To further support this finding, we utilized the complementary approach of nuclear protein fractionation, which revealed an increase in cytosolic FOXO1 and a concomitant decrease in nuclear FOXO1 in TAZ-KO cells upon inhibition of AMPK (Fig. 6B).

**Figure 6.**
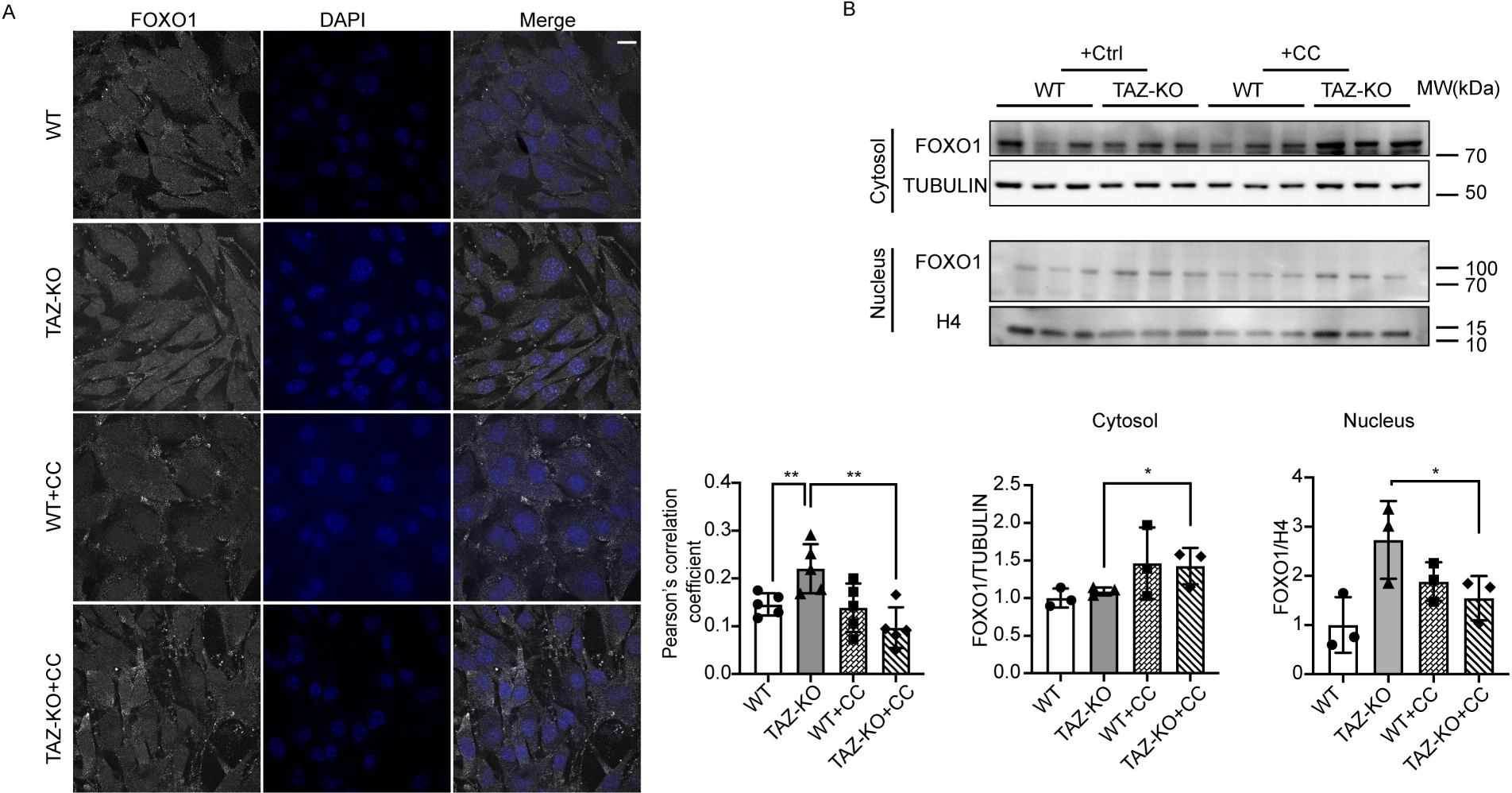
AMPK activation promotes FOXO1 nuclear translocation in TAZ-KO cells. (A) IF staining was used to show the localization of FOXO1 in myoblasts following treatment with CC (5 μM, 16 h) or vehicle. Scale bar = 10 μm. Pearson’s colocalization analysis between DAPI and FOXO1 (bottom); n = 5. (B) FOXO1 protein levels were assayed in nuclear and cytoplasmic cellular fractions following treatment with CC (5 μM, 16 h) or vehicle. Tubulin and histone H3 were used as internal controls for cytoplasmic and nuclear fractions, respectively. Corresponding quantitative analyses are also presented (bottom right). Data points represent mean ±S.D. (*error bars*) for each individual biological replicate of each group. * 0.05<*p*<0.1, ** 0.01<*p*<0.05, *** 0.001<*p*<0.01.

### Glucose uptake is increased in TAZ-KO cells

Given the role of AMPK in regulating glucose metabolism and our data indicating that AMPK is hyperactivated in TAZ-KO cells, we sought to test whether TAZ-KO cells also exhibit altered glucose homeostasis, as reported in BTHS patients (Cade et al., 2019; Cade et al., 2021). Previous studies have shown that pharmacological activation of AMPK using the AMP mimetic, 5-aminoimidazole-4-carboxamide-1-β-o-ribofuranoside (AICAR), results in enrichment of glucose transporter type 4 (GLUT4) in the plasma membrane and a concomitant increase in glucose uptake (Merrill et al., 1997; Kurth-Kraczek et al., 1999). Consistent with this, we observed elevated levels of GLUT4 in TAZ-KO myoblasts, accompanied by an increase in both glucose uptake and cellular glucose concentration (Fig. 7A, B, C).

**Figure 7.**
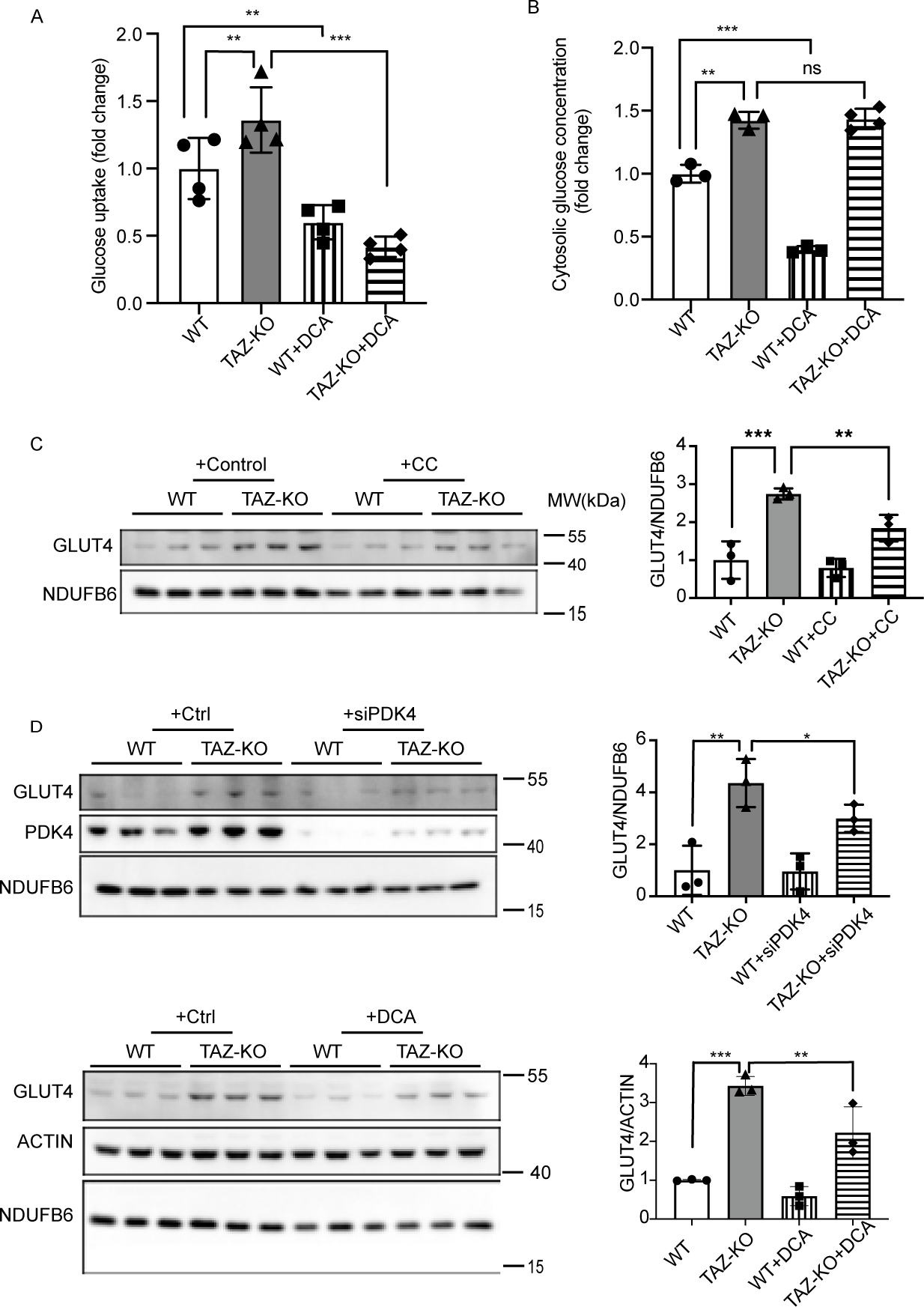
Glucose utilization is dysregulated in TAZ-KO cells. (A) Myoblasts were incubated with 1 mM 2DG. The glucose uptake rate was determined by measuring the amount of 2DG taken up by the cells, with values normalized to total protein in each sample before analysis. (B) Cytosolic glucose concentration was measured according to “Experimental procedures”. (C) (D) GLUT4 protein expression was measured by WB in myoblasts treated with either 5 mM DCA for 24 h or PDK4-targeted siRNA. NDUFB6 and ACTIN were used as internal controls. The data points represent means ±S.D. (*error bars*) for each individual biological replicate of each group. * 0.05<*p*<0.1, ** 0.01<*p*<0.05, *** 0.001<*p*<0.01.

PDH is regarded as a key metabolic regulator capable of altering glucose utilization, though the details of this mechanism are largely unknown. Therefore, we sought to test whether elevated GLUT4 expression is associated with increased PDH activity. To test this, we assayed GLUT4 expression in TAZ-KO cells following inhibition of PDK4 via DCA or siRNA, and observed a concomitant decrease in GLUT4 expression (Fig. 7D). To establish whether regulation GLUT4 by PDK4 is dependent on AMPK activation in TAZ-KO cells, we knocked down PDK4 and assayed levels of pAMPK and FOXO1. Surprisingly, PDK4 knockdown resulted in reduced levels of pAMPK and FOXO1 (Fig. 8), suggesting that a positive feedback loop exists between upregulation of PDK4 and activation of AMPK.

**Figure 8.**
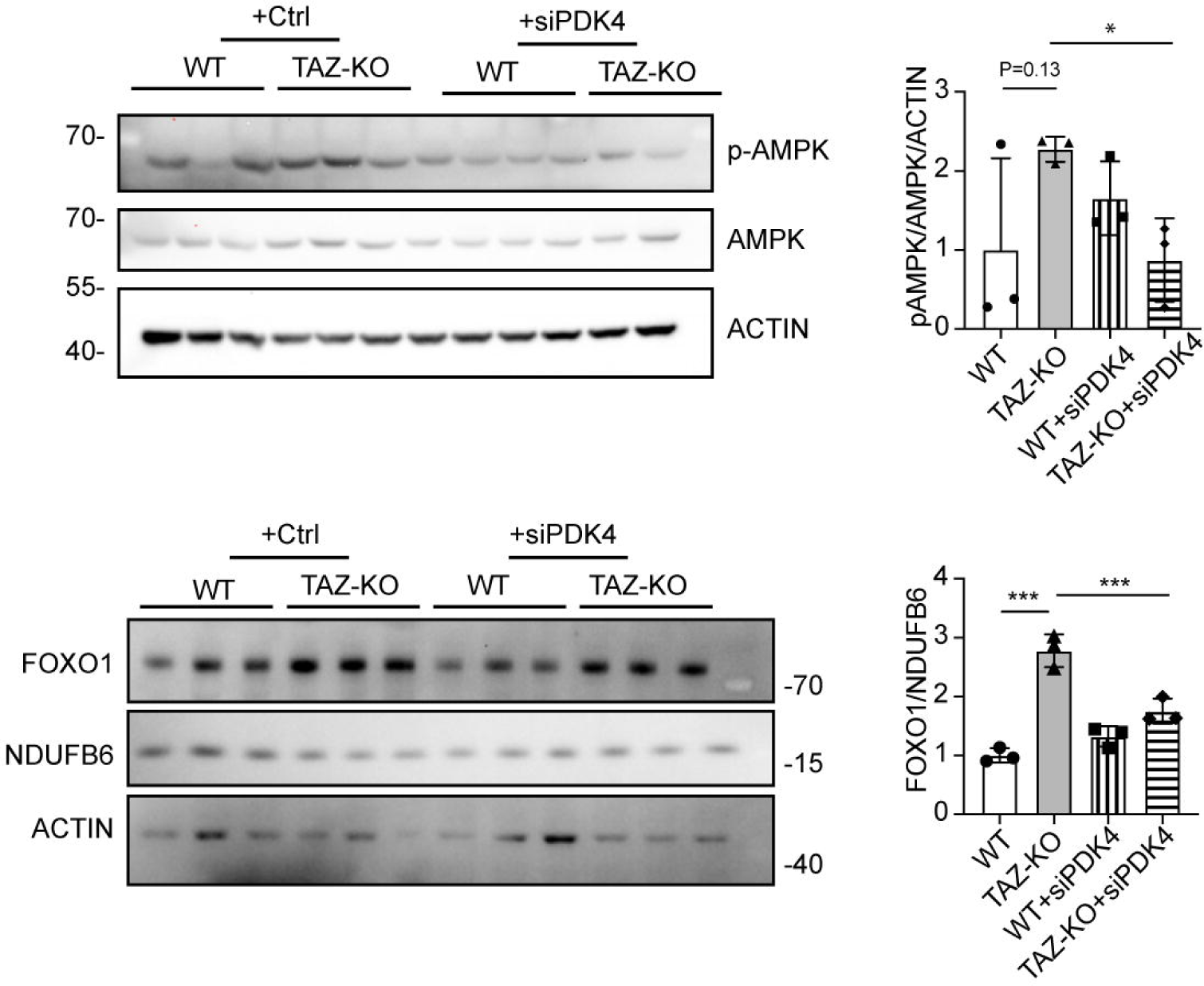
PDK4 upregulation in TAZ-KO cells exacerbates AMPK activation and FOXO1 expression. Phosphorylation of AMPK residue Thr^172^ (p-AMPK) and FOXO1 protein levels were assayed in WT and TAZ-KO myoblasts treated with PDK4-targeted siRNA. The data points represent means ±S.D. (*error bars*) for each individual biological replicate of each group. * 0.05<*p*<0.1, *** 0.001<*p*<0.01.

Taken together, these data support the hypothesis that, in tafazzin-deficient cells, hyperactivation of AMPK induces nuclear translocation of FOXO1, leading to upregulation of PDK4, which potently inhibits PDH activity. Activation of AMPK and upregulation of PDK4 also induced changes in glucose homeostasis by increasing glucose uptake and intracellular glucose levels, presumably through increased expression of the glucose transporter GLUT4.

## Discussion

This study delineates the mechanistic basis for PDK4 upregulation in TAZ-KO cells, leading to potent inhibition of PDH activity. Based on this model, metabolic stress associated with tafazzin deficiency causes hyperactivation of AMPK. Activated AMPK phosphorylates FOXO1, inducing its nuclear translocation and robust transcriptional upregulation of PDK4. Upregulation of PDK4 in tafazzin-deficient cells causes direct phospho-inhibition of PDH activity, accompanied by elevated expression of the glucose transporter GLUT4, increased glucose uptake, and activation of a positive feedback loop to enhance AMPK signaling (Schema 1).

**Schema 1.**
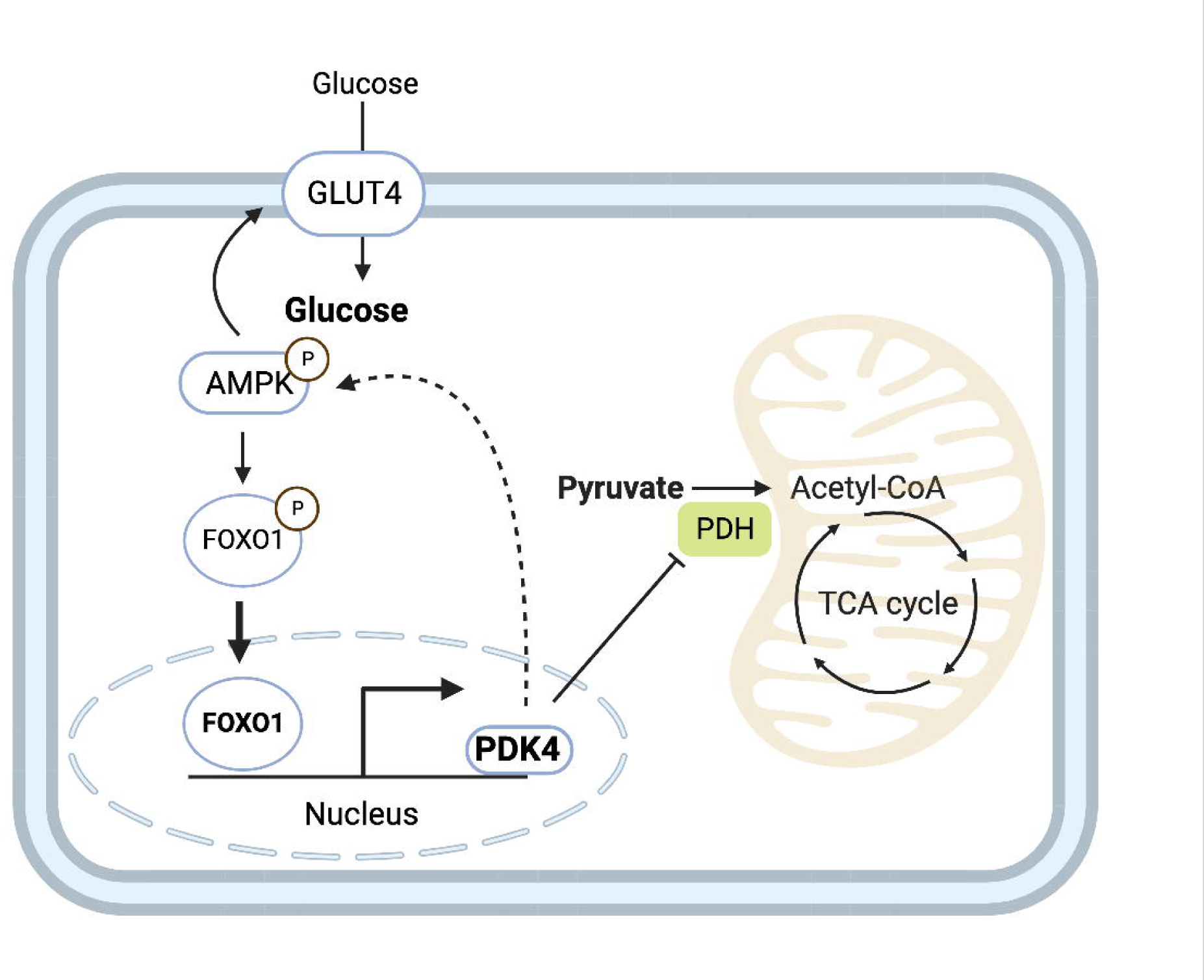
Proposed mechanism underlying metabolic perturbation in TAZ-KO cells. Activation of AMPK in TAZ-KO cells leads to enrichment of FOXO1 in the nucleus, where it acts to increase transcription of *Pdk4*. Upregulation of PDK4 results in altered cellular metabolism by inactivating PDH and applying positive feedback to activate AMPK. This mechanism represents a potential link between tafazzin-deficiency and the metabolic alterations observed in BTHS.

The observation of PDH phospho-inhibition is consistent with previous reports (Li et al., 2019; Greenwell et al., 2021). Li et al. showed that acetyl-CoA levels are substantially reduced in TAZ-KO myoblasts as a result of PDH inhibition. Acetyl-CoA can be produced through multiple pathways, including through the PDH-mediated conversion of glycolysis-derived pyruvate and the metabolic breakdown of fatty acids during FAO.

In the current study, we provide evidence supporting the upregulation of glucose uptake and the accumulation of intracellular glucose in TAZ-KO cells. Our findings suggest that intracellular glucose accumulates due to AMPK activation, which is consistent with what has been previously reported in myocardial extracts from tafazzin-knockdown mice (Greenwell et al., 2021). Upregulation of PDK4 is a sensitive indicator of increased FAO (Rowles et al., 1996; Pettersen et al., 2019), and AMPK activation has been previously linked to membrane enrichment of the fatty acid translocase CD36 (Ren et al., 2016) and a reduction in malonyl-CoA-mediated inhibition of mitochondrial fatty acid uptake (Hopkins et al., 2003). Surprisingly, we observed no significant increase in FAO in TAZ-KO cells compared to WT controls (Fig. 9). Although FAO represents a more substrate-efficient means of producing energy, compensatory upregulation of energy production via FAO is likely hindered in TAZ-KO cells by the previously reported deficiencies in TCA cycle function and ETC complex stability and/or problems with import/processing of fatty acids in the mitochondrial inner membrane (Kessler and Friedman, 1998; Pfeiffer et al., 2003; McKenzie et al., 2006; Li et al., 2019). Thus, the inability of TAZ-KO cells to meet their energetic demands as a result of deficient TCA and ETC function may serve as a metabolic trigger for activating the stress sensor AMPK and promoting its downstream effects on the FOXO1-PDK4-PDH pathway.

**Figure 9.**
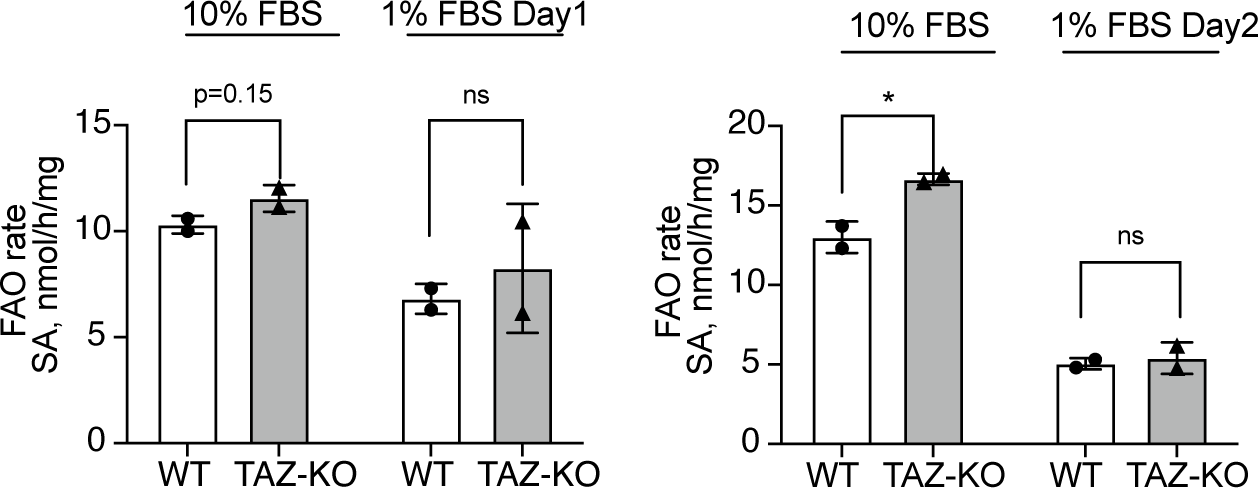
Fatty acid oxidation is not significantly changed in TAZ-KO cells. WT and TAZ-KO cells were incubated with tritium-labeled oleate to allow for import and subsequent oxidation. FAO rate was calculated based on the amount of tritium-labeled H^+^ released back into the medium as a result of oleate oxidation; n = 4. Data points represent means ±S.D. (*error bars*) for each individual biological replicate of each group. * 0.05<*p*<0.1, ** 0.01<*p*<0.05, *** 0.001<*p*<0.01.

Greenwell et al. observed a similar reduction in PDH activity in myocardial extracts from tafazzin-knockdown mice compared to WT littermates (Greenwell et al., 2021). Interestingly however, they reported no significant difference in the phosphorylation profile of PDH between knockdown and WT mice and also noted a reduction in PDK4 mRNA and no change in FOXO1 mRNA levels. These findings are difficult to reconcile with the current study, but they highlight the possibility that the exact mechanism by which tafazzin deficiency inhibits PDH activity may exhibit tissue specificity.

Recent studies have identified similar abnormalities in metabolic substrate utilization associated with tafazzin deficiency. For example, Cade and coworkers reported elevated glucose turnover rate and reduced fatty acid utilization in serum samples from patients with BTHS relative to healthy controls, whereas in myocardial tissue, they observed no significant difference in FAO but a nearly 2-fold increase in glucose oxidation (Cade et al., 2019; Cade et al., 2021). More importantly, it was found that alterations in substrate oxidation are positively corelated with cardiac dysfunction in BTHS patients, underscoring the importance of understanding the associated regulatory pathways (Cade et al., 2021). In contrast to the findings from patients, recent work by Greenwell and colleagues provided evidence of impaired glucose oxidation and elevated FAO in isolated hearts from tafazzin-knockdown mice (Greenwell et al., 2021). While these findings differ slightly from each other and the current study, it is pertinent to note that all three lines of research have independently identified dysregulation of metabolic substrate utilization as a conserved outcome associated with tafazzin deficiency, highlighting the importance of follow-up studies in additional BTHS models (e.g., patient-derived lymphoblasts) to clarify the mechanistic differences.

Our findings are the first reported link between the transcription factor FOXO1 and the regulation of metabolic homeostasis in the context of BTHS. Intriguingly, numerous studies have implicated FOXO1 in the inhibition of MyoD-mediated myogenic differentiation (Khan et al., 1999; Calhabeu et al., 2013; Xu et al., 2017; Milewski et al., 2021), and recent work suggests that suppression of MyoD may underlie BTHS-associated skeletal muscle myopathy (Vo et al., 2023). Therefore, it is tempting to speculate that AMPK-mediated activation of FOXO1 could represent a shared regulatory mechanism linking together the metabolic and muscular pathologies of BTHS.

The goal of the current study was to elucidate the mechanism underlying the suppression of PDH activity in tafazzin-deficient cells. Because PDH acts as a regulatory hub to govern metabolic substrate utilization, and differential substrate oxidation has been linked to cardiac dysfunction in BTHS, these findings may help to guide the development of effective therapies for treating BTHS. For example, previous work has shown that pharmacological inhibition of FOXO1 can ameliorate cardiac defects associated with type 2 diabetes by suppressing PDK4 and thereby increasing PDH activity (Gopal et al., 2021). Direct inhibition of PDK4 with DCA has been previously tested for its ability to improve cardiac function in tafazzin-knockdown mice, and although no significant improvement in cardiac structure or function was observed, DCA treatment was reported to improve both PDH activity and myocardial glucose oxidation (Greenwell et al., 2021). Thus, future studies should be aimed at testing the ability of these drugs and others that target the AMPK-FOXO1-PDK4 pathway to specifically improve the metabolic irregularities associated with BTHS.

### Experimental procedures

#### Cell lines and growth conditions

The TAZ-KO C2C12 mouse myoblast model was generated previously from WT C2C12 cells using CRISPR/Cas9 targeted against exon 3 of the *TAFAZZIN* gene (Lou et al., 2018). Off-target analysis was performed based on the top 10 predicted off-target sites as previously described (Li et al., 2019). Cells were grown in Dulbecco’s modified Eagle’s medium (DMEM; Gibco) containing 4.5 g/L glucose, supplemented with 10% fetal bovine serum (Hyclone), 2 mM glutamine (Gibco), penicillin (100 units/mL), and streptomycin (100 μg/mL). All cell lines were incubated in 5% CO_2_ at 37°C. To induce myoblast differentiation, WT and TAZ-KO C2C12 cells reaching 80% confluency were switched to DMEM containing 2% horse serum (Gibco), 2 mM glutamine (Gibco), penicillin (100 units/mL), and streptomycin (100 μg/mL) (both from Invitrogen Corp.) until the indicated time points. The fresh medium for differentiation induction was changed every 48 h.

#### Mouse heart tissue preparation

The TAZ-KO mouse model was developed in the C57BL/6 mouse background by Dr. Douglas Strathdee (Beatson Institute, United Kingdom) as previously described (Cadalbert et al., 2015). WT and TAZ-KO mice (3 each) were euthanized to harvest hearts and skeletal muscles, and tissues were stored at −80°C. One day before isolation, tissues were preserved in RNAlater™-ICE (Invitrogen) and extracted using an RNeasy kit (QIAGEN). To isolate total protein, 30 – 40 mg of tissue was homogenized by sonication in RIPA buffer with an optimal amount of protease inhibitor (Roche) on ice. After centrifugation, the supernatants containing protein were used for immunoblotting analysis.

#### siRNA construct and incubation

siRNAs targeting PDK4 (SR413673) and scrambled negative control siRNA (SR30004) were purchased from Origene. siRNA was transfected using TransIT-X2® Dynamic Delivery System (Mirus) on the 2^nd^ day after cell seeding and cultured for 24 h before collection.

#### RNA extraction and real-time qPCR analysis

cDNA was synthesized from total RNA using SuperScript™ (Invitrogen). Real-time qPCR was carried out with the QuantSudio3 real-time qPCR machine (Applied Biosystems) using PowerUp™ SYBR™ Green reagent (Thermo Fisher). Relative mRNA transcript levels were quantified with the 2–ΔΔCT method (where CT is threshold cycle), using β-actin as an internal control. Primers used in this study are listed in Table 1.

**Table 1:**
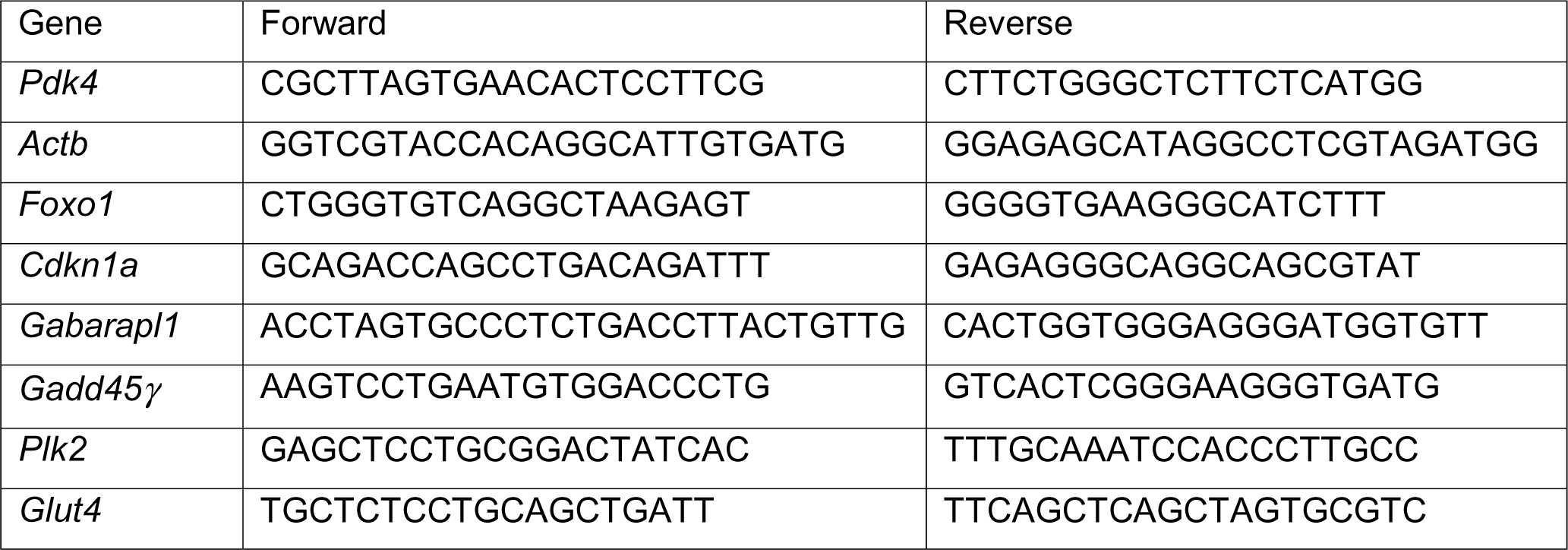
qPCR Primers used in this paper:

#### Western blot analysis

Protein was extracted in RIPA buffer with proteinase and phosphatase inhibitor cocktail (ChemCruz). 20 μg of denatured total protein for each sample was loaded into an 8%-12% SDS-PAGE gel and separated by electrophoresis. The gel was transferred onto polyvinylidene fluoride (PVDF) membranes using a wet transfer method. After transfer, the membranes were probed with the following primary antibodies against PDK4, 1:1000 (ab214938; Abcam); NDUFB6, 1:10000 (ab110244; Abcam); FOXO1, 1:1000 (149527; Cell Signaling); FOXO1, 1:1000 (2880; Cell Signaling); ACTIN, 1:1000 (sc47778; Santa Cruz); p-AMPK (Thr172), 1:2000 (2535S; Cell Signaling); AMPKα 1:1000 (2793S; Cell Signaling); TUBULIN, 1:10000 ab184970; Abcam); H4, 1:1000 (ab16483; Abcam); H3, 1:500 (sc517576; Santa Cruz) GLUT4, 1:2000 (66846; Proteintech); and p-PDH (Ser293), 1:1000 (AP1062; Millipore). Anti-mouse or anti-rabbit secondary antibodies (1:10000) were incubated for 1 h at room temperature, and membranes were developed using SuperSignal™ West Pico PLUS or West Atto ECL (Thermo Scientific).

#### Nuclear protein fractionation

2×10^7^ cells were suspended in lysis buffer 1 (50 mM Tris-HCl pH 7.4,100 mM NaCl, 0.1% NP-40, and freshly added protease inhibitor cocktail). The pellet was collected after centrifugation for 5 m at 1200 x *g* followed by washing in 1 mL of lysis buffer with a serial increase of NP-40 from 0.2% to 0.5%. The pellet was then resuspended in 500 μL of lysis buffer 2 (50 mM Tris-HCl pH 7.4, 100 mM NaCl, 1% NP-40, 0.1% SDS, 0.5% sodium deoxycholate, and freshly added protease inhibitor cocktail). The supernatant was collected and used as the nuclear lysate.

#### Chromatin immunoprecipitation

C2C12 myoblasts (1×10^7^) were harvested and treated with 1% paraformaldehyde (PFA) to cross-link proteins to DNA. The cells were lysed in the presence of protease inhibitors (Roche), sonicated to shear DNA into fragments, and incubated with an antibody against FOXO1 (1:50, 2880; Cell signaling) or normal IgG overnight. The purified DNA and input genomic DNA were analyzed by real-time qPCR. The primer sequences for the PDK4 promoter used in this experiment were as follows: forward 5’-CAA CCA AGT CGT TAC AGC GT −3’ and reverse 5’-TCT TTC TTC ATT GCG GCT GC −3’.

#### Glucose concentration and uptake measurement

Medium: Cells (2000 cells/well) were plated in 96-well cell culture plates in 200 μL of complete DMEM and grown at 37°C for 72 h. At 24 h and 48 h, cells were counted using a SpectraMax i3x plate reader with MiniMax 300 imaging cytometer (Molecular Devices), and 5 μL of medium was taken and further diluted 40-fold with phosphate-buffered saline (PBS). A glucose detection reagent (Glucose-Glo kit, #J6021, Promega) was added to measure glucose concentrations according to the manufacturer’s instructions, and luminescence was detected using a SpectraMax i3x plate reader (Molecular Devices).

Lysate: After 48 h, each well was washed with PBS and lysed using 0.6 N HCL, followed by 1 M Tris base neutralization. Luminescence was detected using a SpectraMax i3x plate reader (Molecular Devices).

Glucose uptake was measured by a commercially available glucose uptake kit (#J1341, Promega) following the manufacturer’s directions with mild alterations. Briefly, 2×10^6^ cells were harvested and washed with PBS. The cells were divided into two groups and treated with either 1 mM 2-Deoxy-D-glucose (2DG) or vehicle control for 30 m at room temperature. Cells were then washed to remove 2DG, sonicated briefly, then added to a 96-well plate. Stop buffer and neutralization buffer were sequentially added. 2DG6P detection reagent was then added, followed by a 1 h incubation at room temperature. Luminescence was detected using a SpectraMax i3x plate reader (Molecular Devices). When applicable, cells were treated with DCA for 24 h before adding 2DG.

#### Fatty acid oxidation rate

LC-FAO flux was determined by measuring the production of radiolabeled H_2_O from [9,10-^3^H(N)]-oleic acid, as described previously (Manning et al., 1990; Olpin et al., 1997; Olpin et al., 1999). Measurements were made at 37 °C in triplicate, and oxidation rates were expressed as nanomoles of fatty acid oxidized per hour per milligram of cellular protein.

#### Immunofluorescence

WT and TAZ-KO C2C12 cells were grown on coverslips in 24-well dishes. The cells were washed three times with PBS and then fixed in 4% PFA + 8% sucrose (3:1) at 37°C for 20 m. After fixation, the cells were washed twice with PBS and then permeabilized in PBS containing 0.3% TritonX100. The cells were washed with PBS three times, then incubated in blocking solution (5% BSA diluted in PBS) at room temperature for 1 h before incubating with FOXO1 antibody (#2880, Cell Signaling; diluted 1:150 in 5% BSA/ sc374427, Santa Cruz; 1:100) at 4°C for a minimum of 16 h. Cells were then washed three times with PBS before being incubated with Alexa Fluor 488 conjugated secondary antibody (A32723, Thermo Fisher; diluted 1:200 in 5% BSA) at room temperature for 1 h. After three final washes with PBS, the slides were mounted with ProLong™ Diamond Antifade Mountant containing DAPI (Invitrogen). All images were captured using a Leica TCS SP8 confocal microscope.

#### Statistical analysis

All values are presented as means ± SD. Statistical analyses were performed in GraphPad Prism software using a two-tailed unpaired Student’s *t*-test on data obtained from at least three independent experiments, which were performed on different days with different biological replicates.

## CRediT Author Statement

**Z.L.** – conceptualization, methodology, investigation, validation, formal analysis, visualization, writing – original draft; **M.W.S.** – resources, validation, writing – revision, review, & editing; **P.L.** – investigation, validation; **S.D.** – investigation, validation; **M.R.** – resources, validation; **M.S.** – resources, validation, funding acquisition; **T.R.E.** – investigation, validation; **M.B.** – investigation, validation; **M.C.** – investigation, validation; **S.H.** – investigation, validation; **R.H.H.** – methodology, validation, funding acquisition; **M.L.G.** – conceptualization, validation, project administration, funding acquisition, writing – review & editing.

## Acknowledgements

The authors would like to thank Dr. Peter G. Barth (Amsterdam UMC), Dr. Peter W. Stacpoole (University of Florida), Dr. Charles E. McCall (Wake Forest University); and Dr. Vishal Gohil (Texas A&M University) for comments on the manuscript.

